# SmartRNASeqCaller: improving germline variant calling from RNAseq

**DOI:** 10.1101/684993

**Authors:** Mattia Bosio, Alfonso Valencia, Salvador Capella-Gutierrez

## Abstract

**Background:** Transcriptomics data, often referred as RNA-Seq, are increasingly being adopted in clinical practice due to the opportunity to answer several questions with the same data - e.g. gene expression, splicing, allele-specific expression even without matching DNA. Indeed, recent studies showed how RNA-Seq can contribute to decipher the impact of germline variants. These efforts allowed to dramatically improved the diagnostic yield in specific rare disease patient cohorts. Nevertheless, RNA-Seq is not routinely adopted for germline variant calling in the clinic. This is mostly due to a combination of technical noise and biological processes that affect the reliability of results, and are difficult to reduce using standard filtering strategies.

**Results:** To provide reliable germline variant calling from RNA-Seq for clinical use, such as for mendelian diseases diagnosis, we developed SmartRNASeqCaller: a Machine Learning system focused to reduce the burden of false positive calls from RNA-Seq. Thanks to the availability of large amount of high quality data, we could comprehensively train SmartRNASeqCaller using a suitable features set to characterize each potential variant.

The model integrates information from multiple sources, capturing variant-specific characteristics, contextual information, and external sources of annotation. We tested our tool against state-of-the-art workflows on a set of 376 independent validation samples from GIAB, Neuromics, and GTEx consortia. SmartRNASeqCaller remarkably increases precision of RNA-Seq germline variant calls, reducing the false positive burden by 50% without strong impact on sensitivity. This translates to an average precision increase of 20.9%, showing a consistent effect on samples from different origins and characteristics.

**Conclusions:** SmartRNASeqCaller shows that a general strategy adopted in different areas of applied machine learning can be exploited to improve variant calling. Switching from a naïve hard-filtering schema to a more powerful, data-driven solution enabled a qualitative and quantitative improvement in terms of precision/recall performances. This is key for the intended use of SmartRNASeqCaller within clinical settings to identify disease-causing variants.

## Background

Being able to associate genomic variation to phenotypic traits is a long-lasting question and fundamental task for omics data analysis. Massive adoption of next sequencing technologies enabled the discovery of causal links between genetic variants and phenotypes. This is especially true for monogenic mendelian diseases (1, 2) and in most of cancer studies (3–5). On one side, NGS data have been used to elucidate the genetic origin of many diseases, with successful diagnoses in 41% of cases overall. On the other side, hundreds of cancer driver genes, and thousands of putative cancer-driver mutations have been identified using NGS with important consequences for diagnosis and treatment.

Whole-genome sequencing (WGS) and whole-exome sequencing (WES) are commonly adopted both in multicenter studies with thousands of patients (6–8), and increasingly in clinical daily practice (2, 9–11). In parallel, initiatives like GTEx (8) showed how RNA-Seq data enriched the picture of genome-phenome relationships, for example defining tissue-specific expression and eQTLs. The potential to answer multiple questions simultaneously from RNA-Seq e.g. gene expression, splicing detection, allele specific expression (12–15), jointly with its reduced costs, convinced an increasingly large share of scientists to adopt RNA-Seq in their analyses.

Using RNA-Seq to call germline variants can be beneficial in clinical settings, for example for Mendelian and common diseases studies. While RNA-Seq does not require additional laboratory experiments if data are already collected, it can enhance the information from samples without matching DNA (16, 17). Indeed, it has been shown to significantly improve the diagnostic yield for Rare Diseases (18) when used jointly with DNA data, and thoroughly processed by field-experts. These recent results show an opportunity to develop tools to automatically enhance the information that can be extracted from an ever growing number of RNA-Seq samples. Such tools need to deal with a whole set of technical challenges i.e. split read mapping, alternative splicing, RNA-Edit, RNA polymerase errors during transcription, and allele specific expression (12, 15, 16) hindering the reliability of RNA-Seq variant calls. A fundamental step for a broader RNA-Seq adoption in clinical settings for variant discovery and prioritization is to reduce the burden of false positive calls. A number of workflows have been developed to reliably call and filter germline variants from RNA-Seq including SNPiR, Opossum or eSNV-detect (16, 19, 20). Those workflows rely on a set of hard-filtering rules implying a trade-off between quality and quantity of called variants. Such filtering schemas have a limited ability to capture complex patterns, and to discriminate true germline calls from the rest.

In this work, we developed SmartRNASeqCaller, a machine-learning module to accurately predict germline variants from RNA-Seq. It makes use of a Random Forest (RF) model that integrates intrinsic variant features with external annotations. SmartRNASeqCaller then generates a data-driven nonlinear predictor for germline variants, harnessing the power to detect complex feature relationship from a massive high-quality training dataset. With SmartRNASeqCaller we aim to improve existing state-of-the-art in discriminating true germline variants from the rest by adopting a more powerful and integrative approach than the hard-filtering strategy used in most of the existing workflows. The overall objective is to minimize the burden of false positive calls from RNA-Seq to call variants with comparable reliability to WGS/WES results. Similar to other biomedical research fields where machine learning techniques are used (21, 22), the main novelty of our approach relies on learning complex patterns to discriminate if a given call is a true germline variant.

SmartRNASeqCaller can be applied as a standalone module to refine the results from previous variant calling workflows without requiring a full sample re-analysis. In this work, we provide SmarRNASeqCaller as a plugin to the GATK best-practices workflow. This module can be easily integrated into any variant calling workflow, as long as it provides an aligned BAM file, and a VCF file with the variants to be classified.

In order to compare the performance of this newly proposed module, we benchmarked the impact of including SmartRNASeqCaller as an additional step after using the GATK best practices workflow against only using the GATK workflow and against SNPiR. We analysed a set of 10 independent high-quality samples from Neuromics consortium (23), as well as on GIAB sample NA12878 (24). We then compared SmartRNASeqCaller impact when applied to the resulting variants from the GATK best practices pipeline on 365 samples from GTEx consortium, collected from 5 tissues from 73 donors. These independent tests serve to confirm the utility of the method in improving germline variant call precision for clinical applications through specific real use-cases.

## Implementation

We have implemented an effective tool to post-process variant calling results from RNA-Seq to reliably identify germline variants. This tool is designed to be used as an additional step in conventional variant calling workflows. It integrates ideas and resources from the literature (12, 13, 19, 25) within a machine learning framework. The driving approach is to use Random Forest (RF), a machine learning technique, to generate a model that is able to discriminate true germline variants from the rest. This process is possible by identifying complex patterns based on variants annotated features coming from multiple sources.

SmartRNASeqCaller is divided in two main steps. First, each variant is annotated with a set of 20 features (table 1). Seven out of them are intrinsic properties including variant type and length, as well as contextual features including external annotations such as the variant in a RepMask region from the UCSC annotation (26), and whether it is annotated into a RNA-Edit site from (25, 27). In parallel, the caller specific features include GATK specific quality values, as well as others such as BaseQRankSum, MQRankSum and ClippingScore. Second, each variant is processed by a classifier that estimates the likelihood of being a true germline variant e.g. appearing in the genomic DNA. Importantly, this classifier model has been generated using a RF approximation, trained on a set of high-quality matched samples of WGS and RNA-Seq with more than 600’000 variants.

**Table 1.** Features considered to train the random forest model.

|  | Name | Selected? | Extra information |
| --- | --- | --- | --- |
| **Intrinsic properties** | Allele ratio | Yes | Alternative allele percentage |
|  | Alt Len | Yes | Length of the alternative allele |
|  | Genotype | Yes | Heterozygous or homozygous call |
|  | DP | Yes | Depth of coverage |
|  | Ref-Alt Len | Yes | Length difference of alternative and reference alleles |
|  | RefLen | No | Length of reference allele |
|  | Type | No | SNP / Indel |
| **Contextual features** | RNA-Edit | Yes | Annotated as RNA-Edit event in databases |
|  | RepMask | Yes | Included in RepeatMasker track from UCSC Genome Browser |
|  | Homopolymer | Yes | Is the variant within a homopolymeric region of genome (5 bases or more) |
|  | SpliceSite | No | Is the variant within 4 nucleotide distances from an exon-intron junction |
| **GATK-specific Annotations** | BaseQRankSum | Yes | Compares the base qualities of the data supporting the reference allele with those supporting any alternate allele. |
|  | ReadPosRankSum | Yes | Tests whether there is evidence of bias in the position of alleles within the reads that support them, between the reference and alternate alleles. |
|  | LikelihoodRankSum | Yes | Compares the likelihood of reads to their best haplotype match, between reads that support the reference allele and those that support the alternate allele. |
|  | ClippingRankSum | No | Tests whether the data supporting the reference allele shows more or less base clipping (hard clips) than those supporting the alternate allele. |
|  | ExcessHet | No | Estimates excess heterozygosity in a population of samples |
|  | MLEAF | No | Maximum likelihood expectation (MLE) for the allele frequency for each ALT allele |
|  | MLEAC | No | Maximum likelihood expectation (MLE) for the allele counts for each ALT allele |
|  | MQ0 | No | Count of all reads that have MAPQ = 0, it can be used for quality control; |
|  | MQRankSum | No | Compares the mapping qualities of the reads supporting the reference allele with those supporting the alternate allele. |

### Samples for the study

To train and validate our tool, we processed samples from three high-quality independent datasets. First, we use 20 samples from Neuromics consortium with high-quality matching DNA sequencing data, specifically WGS from blood samples, and RNA-Seq obtained from skin fibroblast biopsies. For this work purposes, we considered the DNA variant calling results as our reference set of ground truth variants against which measure the RNA-Seq workflows results. This dataset was split into 10 samples for training and 10 for validation guaranteeing the independence of both subsets as we are interested in the general applicability of the model for identifying true germline variants. Second, we analyzed sample NA12878 from the Genome in a Bottle (GIAB) consortium (24). Specifically, we used RNA-Seq reads from SRR1153470 sample and as gold-standard the set of high-confidence SNPs, small indels, and homozygous reference calls associated to GIAB sample NA12878. Third, we used data from 365 GTEx tissue-samples from 73 donors with matching whole blood WGS callsets from GTEx v7 consortium (35). We limited our scope to 5 tissues per donor: Whole blood, Sun Exposed Skin, Adipose Subcutaneous tissue, Skeletal Muscle, and Fibroblasts. We chose these tissues because they represent the most common tissues collected and/or derived in the clinical practice. They are relatively easy to acquire from patients in routine biopsies, and present different expression profiles and transcriptome complexities (28), representing a good testbed for the most common scenarios in which SmartRNASeqCaller could be applied.

### Baseline variant calling workflow

Prior to the application of SmartRNASeqCaller, we processed RNA-Seq from GIAB and Neuromics with GATK RNA-Seq best practices workflow, available at this repository [https://github.com/inab/RDConnect_RNASeq]. This workflow produces two files i) an aligned BAM file, which is obtained with the STAR v2.35a aligner and uses GATK 3.6.0 for subsequent processing steps (24), and ii) a VCF file with the initial set of candidate variants that will be used as input for SmartRNASeqCaller.

GTEx samples were already aligned with TopHat 1.4, thus we used the provided BAM file as input for the variant calling workflow. This difference in the original alignment step represents an opportunity to evaluate the SmartRNASeqCaller performance on data generated following an alternative approach to the one used to train this classifier.

### SmartRNASeqCaller training

We used 665,178 called variants from 10 matched DNA and RNA-Seq samples from the Neuromics Consortium as our training set. The training dataset size allows to build a model for discriminating true germline variants from the rest using a Random Forest (RF) algorithm with sufficient data to reduce potential overfitting to the training set. We chose to use a RF-based algorithm considering the available number of variants in the training set and the need to detect complex patterns without a predefined structure. Other methods like deep learning require at least a spatial data-structure for building a model. Moreover, RF automatically deals with different data types e.g. binaries, qualitative and quantitative, without requiring prior normalization step, and it is robust to class imbalancing (27, 38). Conversely, Support Vector Machine (SVM) and others classical regression models tend to be more sensitive to the classes unbalanced and, in addition, their performances depend on data normalization strategies (38). Finally, a key aspect for choosing RF over other potential options is the robustness of this approximation to overfitting since we want the model to have consistent performances on novel samples.

An initial set of 20 features, generic and GATK specific, were analyzed for training the model (table 1). We employed a recursive feature elimination strategy with 10 fold cross validation applied on the training variants set (as shown in Figure 1A) to select the best feature set for classification. Analysing the results in Figure 1A, we chose 11 features, given that the overall trade-off among average accuracy, accuracy variance, and overfitting potential of the model. With only 11 features, the overall model accuracy is close to the maximum, is quite compact, and is able to generate robust predictions. Importantly, all excluded features fall very close to some selected feature in the tSNE plot in Figure 1B, suggesting that the information content from the excluded features are already provided by other features in the model. The model features, together with the excluded ones are listed in Table 1. We used the R (version 3.5.1) modules RangeR and caret for the model training and evaluation.

**Figure 1:**
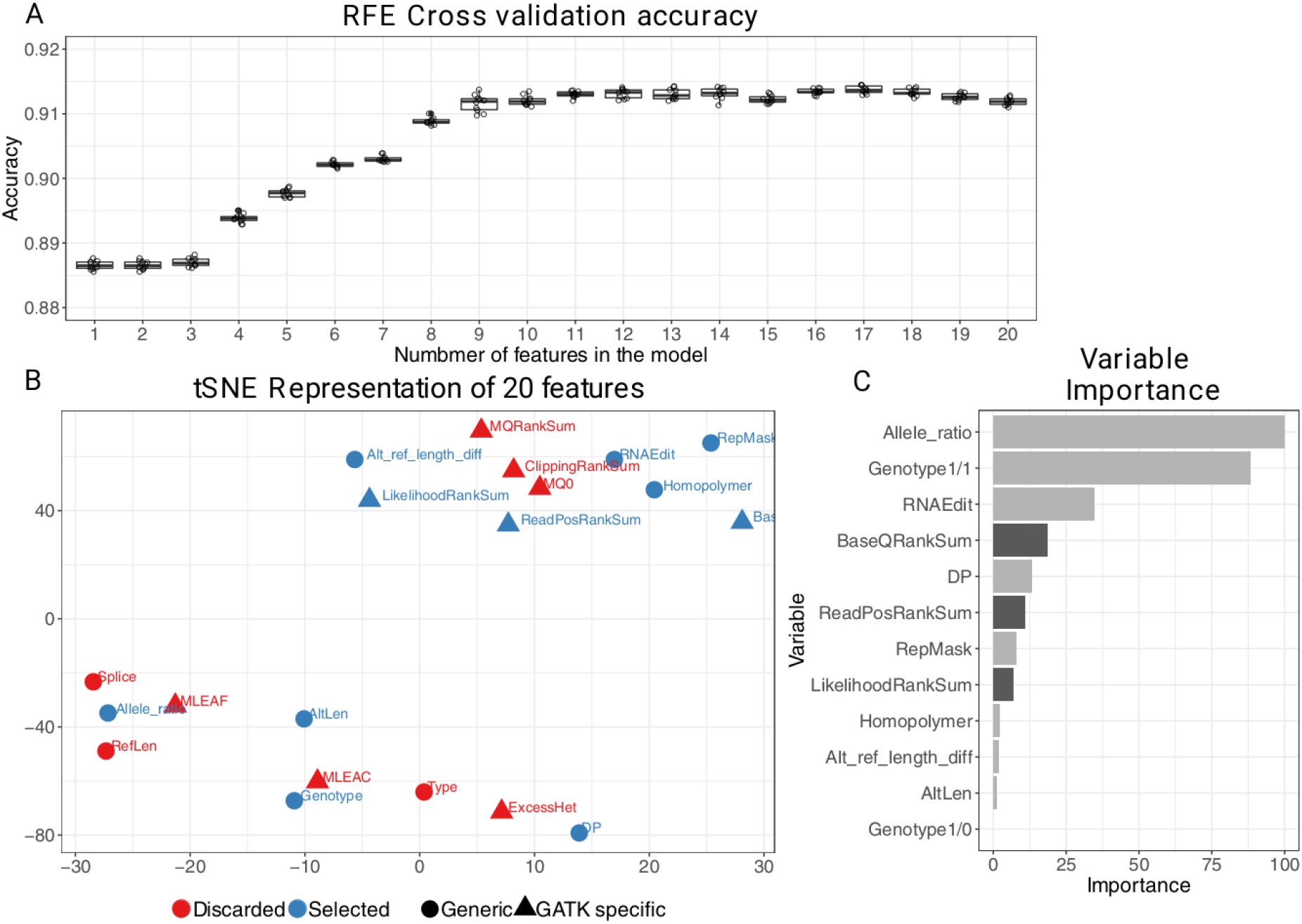
Random forest model construction and iterative feature selection. A) Training performances for recursive feature elimination process. From 11 features on there is no apparent benefit in terms of classification accuracy. B) tSNE representation of the 20 features studied using the training data set. Features are color and shape coded to reflect if they are part of the final model, and if they are generic e.g. intrinsic and contextual properties, or GATK specific. All excluded features are very close to at least one selected one, suggesting that their information content was redundant. C) Variant importance for the prediction model. Light gray darks represent generic annotations, Darker grey bars represent GATK specific annotations.

The selected 11 features are a collection of heterogeneous variant descriptions (Figure 1B and Table 1). It includes intrinsic variant properties as well contextual ones including GATK specific features, the later give an assessment of the trustworthiness of the variant call (table 1). We also included variants annotation from external datasets and genomic context e.g. variant overlapping with an homopolymeric stretch of 5bp or more, variant overlapping with the 4bp intronic region of exon-intron junctions, variant annotated as RNA-Edit events from (16, 34). These external annotations, as remarked in (22), are flags useful to keep or discard a called variant. For instance, SNPiR implemented a series of hard-filtering rules based on those annotations in a subsequent funneling process, progressively reducing the number of potential false positive SNPs in their call set at the cost of strongly reducing the overall number of called variants.

### Model validation

After training the RF model, we tested its predictive performance against 3 other alternative workflows on 10 skin fibroblasts samples from Neuromics, and on sample NA12878. Specifically, we evaluated its predictive performance in terms of precision and recall against the ground truth constituted by genomic high-quality variant calls. Those are the four considered alternatives, including SmartRNAseqCaller.

- GATK Best practices recommendations for calling RNA-Seq variants.
- GATK Best practices recommendations plus SmartRNASeqCaller to validate whether the model refines the initial RNA-Seq called variants.
- SNPiR, which is able to provide reliable calls for SNPs without being limited to somatic variant detection.
- SNPiR-like hard filtering. In this alternative we assess the potential of simple filtering scheme using annotated features for the model. In this workflow we discarded all variants with an annotation of RNA-Edit, homopolymeric region, repmask region, or intron-exon junction. This should serve as a proxy to understand the impact of following a more sophisticated RNA-Seq variant calling approximation. Importantly, this approximation sets the baseline of the performed analysis.

Moreover, we processed 365 samples from GTEx consortium evaluating the impact of including SmartRNASeqCaller on top of GATK best practices workflow. We used the Analysis Freeze WGS variant calls that have been used in GTEx for eQTL and Allele Specific Expression analyses (35) as true reference set. We measured the performances by precision and recall, analyzing the effect both on the bulk of samples and tissue-wise in order to highlight potential biases due to SmartRNASeqCalled being trained on fibroblast samples.

Following commonly accepted practices from genomic data analysis, we focused on regions covered by at least 8 reads. We chose this threshold as it should allow reliable identification for heterozygous genotypes with sufficient sensitivity (22). All samples have been processed using the human reference genome hs37d5 (37).

### Code availability and execution requirements

SmartRNASeqcaller is available at https://github.com/inab/SmartRNASeqCaller. It can be downloaded and executed as a shell script with specific parameters to change its default behaviour, and/or using software containers e.g. dockers, inside a nextflow workflow (29). We expect to guarantee full analysis reproducibility following recommendations around Open Science, Open Data and Open Source. An average run of SmartRNASeqCaller with Nextflow implementation takes 46 minutes, using less than 4 GB RAM with 4 CPUs in parallel.

## Results

Our first goal was to train a reliable model to classify true germline variants using RNA-Seq. Then we validated using three different independent datasets against three commonly used workflows. As demonstrated below, SmartRNASeqCaller would enable the use of RNA-Seq variant calling in the clinic practice by reducing the burden of false positive calls.

### SmartRNASeqcaler obtains better precision/recall results than state-of-the-art workflows on fibroblast samples

We proceeded to measure the SmartRNASeqCaller performance on variants from 10 independent samples from the same Neuromics cohort used for training. We used SmartRNASeqCaller as predictor for all variants considering called variants using WGS as the gold standard. Following broadly adopted practices (19, 19, 30), we evaluated single nucleotide variants in regions with a minimum coverage of 8 or more RNA-Seq reads to reduce the impact of wrong calls due to the effect of random noise on low-coverage areas.

We report the precision/recall results for the all available samples (10 for training set and 10 for validation set) in Figure 2. In the case of SmartRNASeqCaller we reported separately the performance for the training and validation data sets to assess the model robustness and identify potential signs of overfitting.

**Figure 2:**
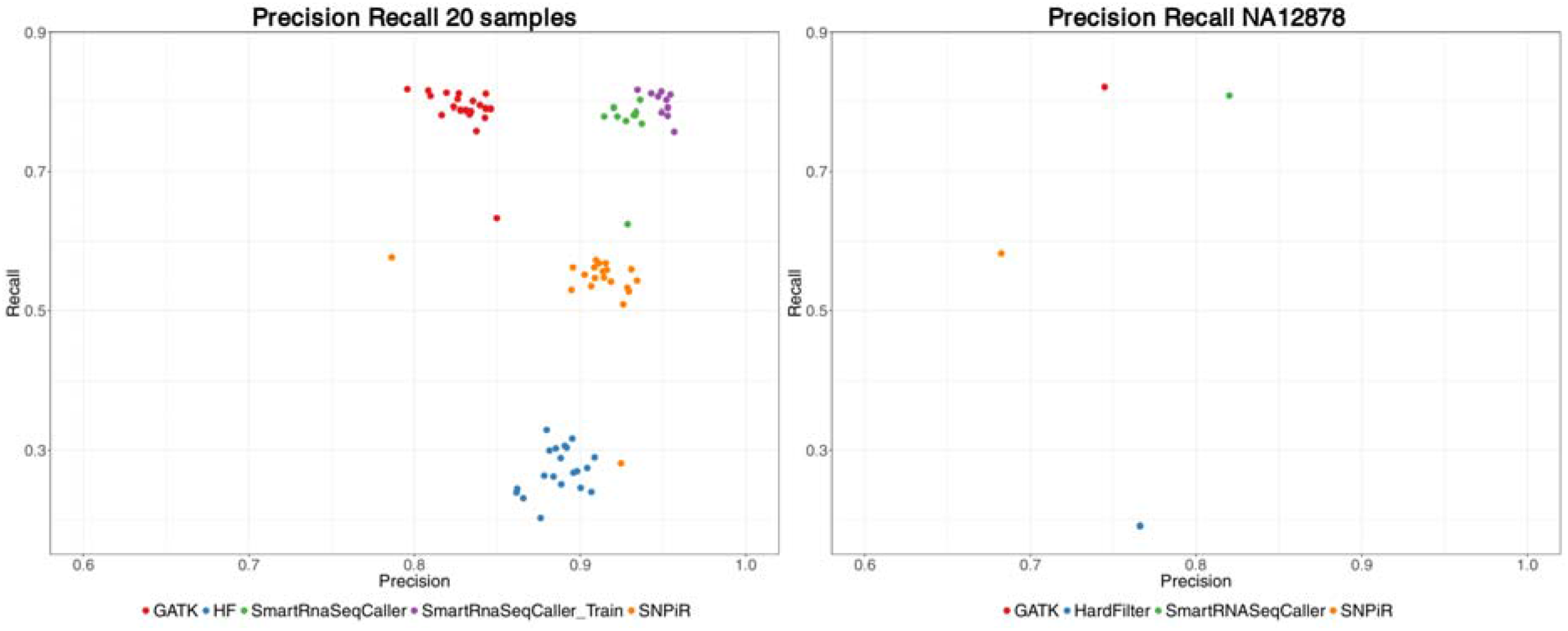
Precision/Recall results on Neuromics and NA12878 GIAB samples. A) Precision/recall results analysing two separated sets of 10 samples each from the same cohort from the Neuromics consortium, which are available at the RD-Connect platform. It compares the SmartRNASeqCaller application against the baseline **GATK** Best practices variant calling workflow, against an alternative naive filtering method depicted as **Hard Filtering**, and against **SNPiR**. We report the results for the training and validation samples for SmartRNASeqCaller separately to show that there is not sign of overfitting to the model. Moreover, we can observe how the strong improvement of precision at a moderate loss of recall behavior is conserved for the validation set of samples, which have not been used at all for generating the Random Forest model As expected, the precision/recall values for the samples in the training set are better than for the validation samples, but the overall effect is similar and robust on the 10 validation samples. Indeed, for the training data set SmartRNASeqCaller achieves +12.0% precision, and 0.1% less recall while that for the validation data set it obtains a +9.3% precision and 0.9% less recall compared to the GATK best practices workflow (supplementary figure 1). B) Precision/recall results after analysing sample NA12878. It compares SmartRNASeqCaller against the baseline GATK best practices variant calling workflow, against an alternative naive filtering method depicted as **Hard Filtering**, and against “SNPiR”. We can observe how the strong improvement of precision at a moderate loss of recall behavior is conserved in this independent sample as well. Here too, the overall relationships among method are conserved, confirming the results previously obtained on the 20 samples from the Neuromics Consortium.

First, the GATK Best practices workflow has an overall good performance in terms of average precision (82.9% ± 3.9%) and recall (78.7% ± 1.4%). Second, the GATK workflow has a better performance than SNPiR for the whole data set when considering average precision and recall with F1 measure (GATK: 0.81 vs SNPiR: 0.66 From Table 2). Third, when comparing the performance on the training and validation samples for SmartRNASeqCaller we can observe that the model is robust to overfitting. The average performance on the training set, albeit better, is not drastically different when compared to the validation samples. Focusing on differential changes with respect to the baseline established by the GATK best practices workflow (Suppl fig. 1), the overall impact of SmartRNASeqCaller brings significant improvements in precision (on average +9% for the validation set) with a modest tradeoff in recall (on average −0.9% for the validation set). This pattern is observed consistently among training and validation samples. Finally, when compared to naïve hard-filtering strategies, we can appreciate that the average precision is marginally improved but the average recall drastically drops, showing how naïve approaches end-up doing more harm than good. These results support the idea of integrating complex patterns derived from different sources, rather than limiting to simpler intersection or union operations, using strategies based on machine learning techniques

**Table 2.** Summary of F1 statistic on Neuromics samples

| F1<br>measure | Hard<br>Filtering | GATK | SmartRNASeqCaller<br>Train | SmartRNASeqCaller<br>Test | SNPiR |
| --- | --- | --- | --- | --- | --- |
| Mean | 0.41 | 0.81 | 0.87 | 0.84 | 0.66 |
| Median | 0.41 | 0.81 | 0.87 | 0.85 | 0.68 |
| Standard<br>Deviation | 0.04 | 0.02 | 0.01 | 0.03 | 0.08 |
| Minimum | 0.33 | 0.73 | 0.85 | 0.75 | 0.43 |
| Maximum | 0.48 | 0.83 | 0.88 | 0.86 | 0.70 |

F1 measure(geometric mean of precision and recall on 20 samples from a cohort from the Neuromics Consortium. From this summary statistic we can infer how the baseline GATK variant calling workflow achieves better results than the simple hard filtering strategy, and the SNPiR algorithm as well. The application of SmartRNASeqCaller to the GATK best practices workflow, allows to further improve the F1 results. We split train and validation values for SmartRNASeqCaller, to avoid bias of the training samples on the overall result.

### SmartRNASeqCaller improves precision on sample NA1278

As a further evaluation step to study the model generalization and to exclude specific biases from the considered samples, we tested SmartRNASeqCaller on the publicly available sample NA12878 from the GIAB Consortium. On one hand, we processed raw RNA-Seq reads through the GATK best practices variant calling workflow to have a baseline calls set. Building on this set we applied SmartRNASeq as an additional step to the GATK Best practices called variants for comparison against it, against SNPiR, and against a naïve hard-filtering strategy. We used GIAB calls from DNA sequencing as the ground truth to evaluate the RNA-Seq variant calling results.

Similarly to the previous analysis, in Figure 2B we reported the performance in terms of precision/recall obtained for SmartRNASeqCaller and other alternative approaches. Similar results to the previously analysed 20 samples were obtained confirming the general usability of our model. Importantly, the baseline established by the GATK best practices workflow yielded better results than SNPiR. This brings in the discussion the impact of previous steps e.g. choice of the alignment strategy as well as the impact of the continuous improvement of external annotation sources.

Similarly to comparison for the Neuromics samples, the application of SmartRNASeqCaller to the baseline results allows to significantly improve precision (8%) with a moderate trade-off in recall (~2%) achieving the best overall results, while the naïve hard-filtering strategy confirms to be the worst performing algorithm due to its drastic effect on the final recall of variants. The baseline values of precision/recall for NA12878 are worse than the average values with Neuromics samples as absolute values. Nevertheless, the change brought by SmartRNASeqCaller is robust and in the same direction, showing how the model behaves consistently across different initial conditions.

### SmartRNASeqCaller is robust to both tissue-of-origin differences, and alignment algorithm

We then assessed SmartRNASeqCaller performance on a large independent cohort from 365 GTEx (8) samples with matching WGS data. We chose tissue from 5 tissues that represent most biopsies in clinical settings: Whole Blood, Skin Sun Exposed, Adipose Subcutaneous, Skeletal Muscle, and Fibroblasts. These tissues have diverse transcriptome complexity and may be a closer representation of datasets used for clinical applications.

GTEx v7 data have been aligned using TopHat v1.4, rather than STAR v3.5.1, which we used to align the training set for SmartRNASeqCaller. Thanks to this, we could test how robust SmartRNASeqCaller is to alternative upstream workflows, as aligners present systematic differences between them. This is a particularly challenging dataset since TopHat 1.4 has been shown to have many limitations and artifacts when compared to recent aligners like STAR or Hisat2 (12, 31).

In Figure 3A, precision/recall results comparing the performance of the baseline TopHat+GATK workflow and SmartRNASeqCaller applied as an additional step to the baseline TopHat+GATK workflow are presented. The overall effect of strong precision improvement with small sensitivity loss observed in Figure 2 is maintained on GTEx data. Indeed, SmartRNASeqCaller improves precision on average by 20.9%, a 6.25 fold greater than the reduction in recall (3.2%).

**Figure 3:**
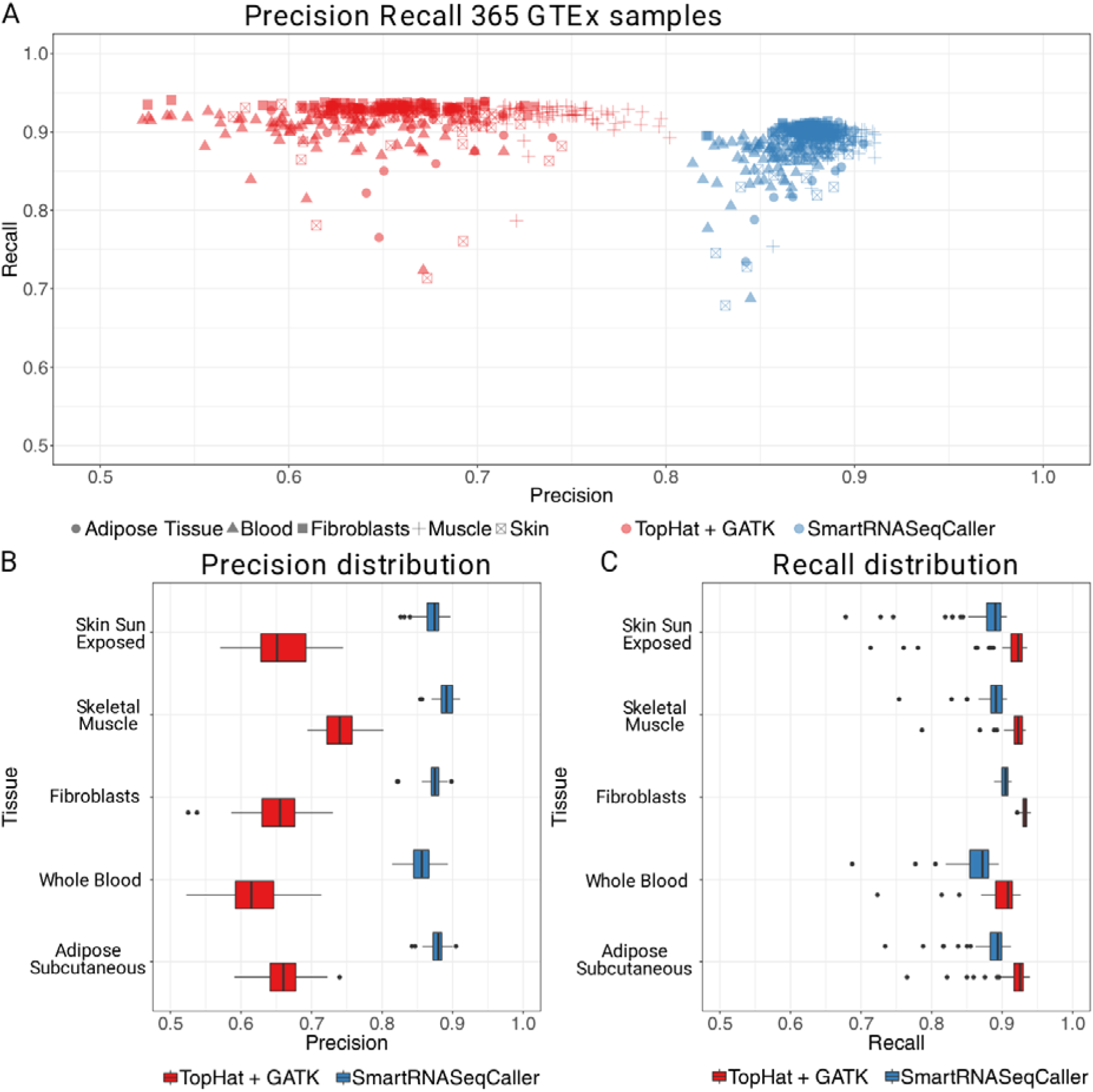
Precision/Recall on 365 GTEx samples. A) Precision/recall results analysing 365 tissue samples from GTEx cohort. Samples are from five different tissues and 73 patients. Precision/recall plot comparing TopHat+GATK Best practices variant calling workflow against SmartRNASeqCaller applied on TopHat+GATK results. SmartRNASeqCaller shows a strong effect improving precision on average by 20.9%, reducing Recall by 3.2% on average. B) Boxplots comparing precision values for GATK best practices and SmarRNASeqCaller. We observe how samples from different tissues have different burden of false positives in GATK. After SmartRNASeqCaller application the differencesare less evident and the boxplots overlap across tissues. C) Boxplots comparing Recall values for GATK best practices and SmarRNASeqCaller. On average, the application of SmartRNASeqCaller reduces recall by 3.2%. The impact of SmartRNASeqCaller is to increase the overall precision, levelling the performance across tissues close to 90%, simultaneously keeping high levels of Recall (between 85 and 90%).

In Figure 3B, we present the precision values separated by tissue and workflow. The median precision values for the TopHat+GATK workflow strongly depend on the tissue of origin, ranging from 61.4% for Whole Blood, to 73.9% for Skeletal Muscle. After the application of SmartRNASeqCaller, the precision levels range increase and are more compact ranging from 85.6% in Whole Blood to 89.1% in Skeletal Muscle samples, reducing dramatically (~50%) the differences between tissues. Similarly to Figure 3B, we present in Figure 3C recall values for tissue of origin and workflow. SmartRNASeqCaller effect is stable across tissues, reducing the sensitivity on average by 3.2% while keeping the average recall between 85%-90% for all analyzed tissues. This is important because we are able to capture much more true germline variants with higher precision that the standard baseline.

In general terms, SmartRNASeqCaller strongly improves the overall precision of RNA-Seq variant calling with a small cost of sensitivity, even for data generated with different aligners and collected from different tissues in the body demonstrating its general applicability.

## Discussion

In this work we developed SmartRNASeqCaller, a random forest model to reliably discriminate true germline variants from the rest using RNA-Seq. SmartRNASeqCaller combines intrinsic variant characteristics, with external annotation sources in a unique model able to reduce the burden of false positive calls from RNA-Seq.

We trained our model using more than 600’000 variants from 10 high-quality samples with matching WGS data from Neuromics Consortium. We then validated it against a dataset of 10 independent samples from the same cohort, as well as on an independent validation set composed by the broadly used sample NA12878 from the GIAB Consortium (24), and by 365 samples from GTEx consortium (8). In all cases, applying SmartRNASeqCaller significantly reduced the number of false positive calls almost halving the number, without hindering recall e.g. average 0.9% loss in recall for the validation samples from GIAB and Neuromics, and 3.2% on GTEx samples. SmartRNASeqCaller allowed to achieve the best precision/recall performance when compared against state-of-the-art workflows e.g. GATK best practices variant calling workflow and SNPiR (16).

A whole set of technical challenges for the wide adoption of RNA-Seq as a source of data for germline variant calling have been described in the literature i.e. split read mapping, alternative splicing, RNA-Edit, RNA polymerase errors during transcription, and allele specific expression (12, 15, 16). Several tools have now been released to address these task-specific challenges. Examples are tools such as STAR and Histat2 (17, 18), which aim to improve read alignment; or REDITools and DeepRed, which are tools to detect RNA Editing events (19, 20). Resources like REDIPortal and RADAR (25, 27) collect regions with evidence of RNA-Edit activity along the human genome and are a valuable resource to spot potential false positive calls.

However, few workflows have been developed to reliably call and filter germline mutations from RNA-Seq. Those developed though rely on a set of hard-filtering rules implying a trade-off between quality and quantity of selected variants. Some examples are eSNV-detect (21), SNPiR (12), and Opossum (22). eSNV-detect (21) combines multiple aligners to reduce aligner-specific errors prior to the variant calling itself. Once this step is completed, eSNV-detect calls variants using SAMtools (32). However, this practice introduces significant computational costs and questions their use in routinary analysis. SNPiR (12) uses BWA-aln (23) to map spliced reads combined with GATK UnifiedGenotyper (24) to generate an initial set of variant calls, which are then filtered using external annotations about variant characteristics e.g. RNA-Edit site, homopolymer region, repmask site. This filtering allows to improve precision at the cost of reduced sensitivity. Opossum (22) employs a different strategy by preprocessing and filtering RNA-Seq raw data to make it suitable for haplotype-based variant calling with Platypus (25). This strategy renders remarkable results, albeit limited to the easily aligned portion of the genome. Moreover, *a priori* exclusion of all sites prone to RNA-Edit, which include many true germline variant e.g. 25% of RNA-Edit positions in RADAR and REDI-Portal databases are located in exonic areas overlap with documented DNA mutations in GnomAD dataset (26), may limit the use of Opossum into routine clinical practice.

Methods evaluation in most of these works is not standardized and is heavily dependent on the annotations used to determine the scope of analysis e.g. gene definitions, inclusion or exclusion of specific regions/SNP type, publicly available gold standard dataset, etc. There is therefore a need to joint efforts in the community to standardize those efforts including the definition of relevant datasets and metrics.

The main driver to develop SmartRNASeqCaller was to obtain the highest reliability for variants called from RNA-Seq experiments for its use in routine clinical practice. For this we focused on improving the precision of the generated variant calls. We first chose to integrate heterogeneous and non-redundant variants features to generate a rich and complex description of each variant. Tools like SNPiR use a similar approach to apply simple filters to exclude variants if characterized by unreliable features, which improved precision compared to baseline. However, a simple filtering strategy is unable to properly exploit the potential of a rich and complex multidimensional space. It can generate a strong tradeoff between precision and sensitivity that can be detrimental for tasks such as diagnosis. For that, we chose to train a Random Forest classifier on more than 600’000 variants from 10 samples. We chose Random Forests because it has been previously applied in complex scenarios with many training samples, producing remarkable results in terms of precision and robustness including DNA variant calling (21). We then evaluated SmartRNASeqCaller following standard practices of processing independent samples from different studies to ensure the general usability of this model across a wide variety of samples from different tissues, and different upstream alternative workflows to generate the initial calls sets.

Here we show that switching from a naïve hard-filtering schema to a more powerful, data-driven solution enabled a qualitative and quantitative improvement in terms of precision/recall. When compared to a SNPiR-like strategies of filtering all variants annotated by some unreliable characteristic, the drastic reduction in recall does not compensate for the improvement in terms of precision. This effect is mostly due to the improvement and expansion of available annotations since the SNPiR publication, as well as to the quality filtering already implemented in the baseline workflow that removes plenty of unreliable variants from RNA-Seq.

SmartRNASeqCaller builds on existing literature for variant calling using RNA-Seq, improving overall performances and trustworthiness of the obtained results. Nevertheless, as noted in (16, 24), its discovery potential is inherently limited by the nature of RNA-Seq experimental set-ups: there is no hope to detect variants in areas of the genome that are not expressed. Similarly, tissue-specific gene expression can limit the discovery of phenotypic-causing variants as many experiment tend to use easily accessible tissues rather than the affected one. Those accessible tissues might not express the genes of interest for dissecting the genetic causes of the observed phenotype. However, recent results showed that it is possible to obtain reliable mutation profile data of not easy-to-reach tissues from other accessible tissues by generating suitable reprogrammed cells (18). How RNA-Seq data is obtained can also directly affect the sensitivity of our method as nonsense variants can be missed as a result of the nonsense-mediated decay mechanisms (33).

Despite these factors limiting the scope of potential discoveries from RNA-Seq, they can simultaneously be turned into a powerful filter against noise. Provided that the sequenced tissue is relevant for the studied disease, RNA-Seq variants can limit the focus to those genes that actually are being used by the affected cells, as well as inferring if there are “missing genes” e.g. genes that are normally expressed in the tissue that are not present in the experiment when considering reference datasets.

An additional factor contributing towards the divergence between RNA-Seq variants and variants extracted from DNA is the existence of genes in which only one parental allele is expressed (16, 34). Previous work in this direction suggests that only 5% – 10% of human genes are subject to monoallelic gene expression (34), which could account for up to half of the missing recall in our results. Strategies to improve the overall recall will require then restructuring baseline variant calling workflows, specifically about the calling and filtering criteria

Although SmartRNASeqCaller allows to drastically reduce false positives from the analyzed data, similarly to other tools e.g. SNPiR, and approximations, our model may miss to filter variants due to systematic errors in the preceding workflows. Different strategies have been proposed to overcome those systematic errors including merging results from multiple samples to exclude novel recurrent rare variants (34). However, we believe that with a much wider and diverse training dataset, the occurrence of systematic errors can be strongly reduced. Moreover, our model can easily incorporate extra features that may characterize systematic errors e.g. DNA sequence surrounding each variant, in future developments.

A go-to RNA-Seq reliable variant calling workflow like SmartRNASeqCaller can help filtering out genomic variants that may look promising from DNA data analysis but are either not expressed in the tissue of interest, or removed by post-transcriptional modifications, reducing the burden of false positive calls and enhancing the diagnosis potential of these analyses.

Importantly, an additional benefit of reliable RNA-Seq variant calling would allow to detect post-transcriptional RNA-specific variants that are not present at genomic level but could have functional effects by themselves and/or jointly with nearby genomic variants. Accurate variant calling results can help investigating if RNA-Edit, generally not considered as source of disease, may act detrimentally towards the cell. It is theoretically possible to detect RNA-Edit events acting like germline variants for further annotation and interpretation for disease generation (35).

## Conclusions

Despite the limitations of calling genomic variants from RNA-Seq, our work demonstrates improvements in the field of RNA-Seq variant calling to detect germline variants with high precision and recall using appropriate machine learning tools.

SmartRNASeqCaller can be a go-to tool for reliable variant calling from RNA-Seq, with the potential to enhance diagnostic yield and have better disease characterization in the tissue of interest. SmartRNASeqCaller allows to harness information from RNA-Seq and to generate a very precise calls set with good sensitivity. These characteristics are of paramount importance in clinical settings and can provide relevant benefits. RNA-Seq can be used to integrate DNA mutation information with tissue specific results providing an independent source of information to filter and validate disease-causing candidate variants.

Furthermore it can palliate the absence of genomic data for specific samples, presenting a viable way to extract a reliable variant calls, and generate a new knowledge base of RNA mutations. This could allow RNA-Seq samples processing for tissues cohorts in clinic to extract a very precise and context-specific mutational landscape without requiring additional DNA sequencing.

Finally, SmartRNASeqCaller can be used as an additional step of any existing variant calling workflow. This makes possible to even reanalyze existing cohorts with the goal of detecting germline variations without requiring expensive computation.

## Declarations

### Ethics approval and consent to participate

Not applicable. All samples processed in this work come from consortia in which the consent has been explicitly granted for research purposes.

### Consent for publication

Not applicable

### Availability of data and material

**Project name**: SmartRNASeqCaller

**Project home page**: https://github.com/inab/SmartRNASeqCaller

**Operating system(s):** Platform independent

**Programming language**: Python, Bash, Nextflow, R

**Other requirements**: GATK 3.6-0, Samtools, Bcftools, Bedtools, tabix, Python 2.7: (pysam, pandas), R 3.5.0 (caret, ranger). Optional: Docker

**License**: GNU GPLv3

### Datasets availability

- GIAB NA12878 data are available at: https://jimb.stanford.edu/giab-resources
- Neuromics cohort: The data that support the findings of this study are available from Neuromics consortium but restrictions apply to the availability of these data, which were used under license for the current study, and so are not publicly available. Data are however available from the authors upon reasonable request and with permission of Neuromics consortium. https://rd-neuromics.eu/project-welcome/
- GTEx data: The data that support the findings of this study are available from GTEx Consortium but restrictions apply to the availability of these data, which were used under license for the current study, and so are not publicly available. Data are however available from the authors upon reasonable request and with permission of GTEx consortium. https://gtexportal.org/home/datasets

## List of abbreviations

RF: Random Forest
WES: Whole Exome Sequencing
WGS: Whole Genome Sequencing
GATK: Genome Analysis ToolKit
GIAB: Genome In a Bottle
GTEx: Genotype Tissue Expression

## Authors’ contributions

**M.B:** Designed the algorithm, performed training and validation of the data, write the first version of the manuscript. **A.V:** Designed the algorithm and validation strategy, **S.C.G** Designed the algorithm and validation strategy, write the final version of the manuscript. All authors read and approved the final manuscript.

## Acknowledgements

We are grateful to Ana Topf and Steve Laurie for facilitating the access to the data from the Neuromics project. We also thank insightful comments over preliminary versions of this work to Marta Melé and Jennifer Harrow.

## References

1. Clark MM, Stark Z, Farnaes L, Tan TY, White SM, Dimmock D, et al. Meta-analysis of the diagnostic and clinical utility of genome and exome sequencing and chromosomal microarray in children with suspected genetic diseases. Npj Genomic Med. 2018 Jul 9;3(1):16.

2. Nguyen MT, Charlebois K. The clinical utility of whole-exome sequencing in the context of rare diseases – the changing tides of medical practice. Clin Genet. 2015 Oct;88(4):313–9.

3. Rau A, Flister M, Rui H, Auer PL. Exploring drivers of gene expression in the Cancer Genome Atlas. Bioinformatics. 2019 Jan 1;35(1):62–8.

4. Thorsson V, Gibbs DL, Brown SD, Wolf D, Bortone DS, Ou Yang T-H, et al. The Immune Landscape of Cancer. Immunity. 2018 17;48(4):812–830.e14.

5. Bailey MH, Tokheim C, Porta-Pardo E, Sengupta S, Bertrand D, Weerasinghe A, et al. Comprehensive Characterization of Cancer Driver Genes and Mutations. Cell. 2018 Apr 5;173(2):371–385.e18.

6. The 1000 Genomes Project Consortium. A global reference for human genetic variation. Nature. 2015 Oct;526(7571):68–74.

7. The UK10K project identifies rare variants in health and disease. Nature. 2015 Oct 1;526(7571):82–90.

8. The Genotype-Tissue Expression (GTEx) project | Nature Genetics [Internet]. [cited 2018 Dec 3]. Available from: https://www.nature.com/articles/ng.2653

9. Zawati MH, Parry D, Thorogood A, Nguyen MT, Boycott KM, Rosenblatt D, et al. Reporting results from whole-genome and whole-exome sequencing in clinical practice: a proposal for Canada? J Med Genet. 2014 Jan;51(1):68–70.

10. Bosio M, Drechsel O, Rahman R, Muyas F, Rabionet R, Bezdan D, et al. eDiVA— Classification and prioritization of pathogenic variants for clinical diagnostics. Hum Mutat [Internet]. 2019 [cited 2019 May 29];0(0). Available from: https://onlinelibrary.wiley.com/doi/abs/10.1002/humu.23772

11. Byron SA, Van Keuren-Jensen KR, Engelthaler DM, Carpten JD, Craig DW. Translating RNA sequencing into clinical diagnostics: opportunities and challenges. Nat Rev Genet. 2016 May;17(5):257–71.

12. Sahraeian SME, Mohiyuddin M, Sebra R, Tilgner H, Afshar PT, Au KF, et al. Gaining comprehensive biological insight into the transcriptome by performing a broadspectrum RNA-seq analysis. Nat Commun. 2017 05;8(1):59.

13. Xu C. A review of somatic single nucleotide variant calling algorithms for next-generation sequencing data. Comput Struct Biotechnol J. 2018 Jan 1;16:15–24.

14. Conesa A, Madrigal P, Tarazona S, Gomez-Cabrero D, Cervera A, McPherson A, et al. A survey of best practices for RNA-seq data analysis. Genome Biol [Internet]. 2016 [cited 2018 Dec 3];17. Available from: https://www.ncbi.nlm.nih.gov/pmc/articles/PMC4728800/

15. Han Y, Gao S, Muegge K, Zhang W, Zhou B. Advanced Applications of RNA Sequencing and Challenges. Bioinforma Biol Insights. 2015 Nov 15;9(Suppl 1):29–46.

16. Piskol R, Ramaswami G, Li JB. Reliable Identification of Genomic Variants from RNA-Seq Data. Am J Hum Genet. 2013 Oct 3;93(4):641–51.

17. Cummings BB, Marshall JL, Tukiainen T, Lek M, Donkervoort S, Foley AR, et al. Improving genetic diagnosis in Mendelian disease with transcriptome sequencing. Sci Transl Med. 2017 19;9(386).

18. Gonorazky HD, Naumenko S, Ramani AK, Nelakuditi V, Mashouri P, Wang P, et al. Expanding the Boundaries of RNA Sequencing as a Diagnostic Tool for Rare Mendelian Disease. Am J Hum Genet. 2019 Mar 7;104(3):466–83.

19. Oikkonen L, Lise S. Making the most of RNA-seq: Pre-processing sequencing data with Opossum for reliable SNP variant detection. Wellcome Open Res [Internet]. 2017 Mar 17 [cited 2018 Dec 3];2. Available from: https://www.ncbi.nlm.nih.gov/pmc/articles/PMC5322827/

20. Tang X, Baheti S, Shameer K, Thompson KJ, Wills Q, Niu N, et al. The eSNV-detect: a computational system to identify expressed single nucleotide variants from transcriptome sequencing data. Nucleic Acids Res. 2014 Dec 16;42(22):e172.

21. Ho DSW, Schierding W, Wake M, Saffery R, O’Sullivan J. Machine Learning SNP Based Prediction for Precision Medicine. Front Genet [Internet]. 2019 [cited 2019 May 29];10. Available from: https://www.frontiersin.org/articles/10.3389/fgene.2019.00267/full

22. Triantafyllidis AK, Tsanas A. Applications of Machine Learning in Real-Life Digital Health Interventions: Review of the Literature. J Med Internet Res. 2019;21(4):e12286.

23. Lochmüller H, Badowska DM, Thompson R, Knoers NV, Aartsma-Rus A, Gut I, et al. RD-Connect, NeurOmics and EURenOmics: collaborative European initiative for rare diseases. Eur J Hum Genet. 2018 Jun;26(6):778.

24. Zook JM, Chapman B, Wang J, Mittelman D, Hofmann O, Hide W, et al. Integrating human sequence data sets provides a resource of benchmark SNP and indel genotype calls. Nat Biotechnol. 2014 Mar;32(3):246–51.

25. Picardi E, D’Erchia AM, Lo Giudice C, Pesole G. REDIportal: a comprehensive database of A-to-I RNA editing events in humans. Nucleic Acids Res. 2017 04;45(D1):D750–7.

26. Karolchik D, Hinrichs AS, Furey TS, Roskin KM, Sugnet CW, Haussler D, et al. The UCSC Table Browser data retrieval tool. Nucleic Acids Res. 2004 Jan 1;32(Database issue):D493–496.

27. Ramaswami G, Li JB. RADAR: a rigorously annotated database of A-to-I RNA editing. Nucleic Acids Res. 2014 Jan 1;42(D1):D109–13.

28. Melé M, Ferreira PG, Reverter F, DeLuca DS, Monlong J, Sammeth M, et al. The human transcriptome across tissues and individuals. Science. 2015 May 8;348(6235):660–5.

29. Di Tommaso P, Chatzou M, Floden EW, Barja PP, Palumbo E, Notredame C. Nextflow enables reproducible computational workflows. Nat Biotechnol. 2017 Apr 11;35:316–9.

30. Laurie S, Fernandez Callejo M, Marco Sola S, Trotta J, Camps J, Chacón A, et al. From Wet Lab to Variations: Concordance and Speed of Bioinformatics Pipelines for Whole Genome and Whole Exome Sequencing. Hum Mutat. 2016 Dec;37(12):1263–71.

31. Baruzzo G, Hayer KE, Kim EJ, Di Camillo B, FitzGerald GA, Grant GR. Simulation-based comprehensive benchmarking of RNA-seq aligners. Nat Methods. 2017 Feb;14(2):135–9.

32. Li H, Durbin R. Fast and accurate short read alignment with Burrows–Wheeler transform. Bioinformatics. 2009 Jul 15;25(14):1754–60.

33. Behm-Ansmant I, Kashima I, Rehwinkel J, Saulière J, Wittkopp N, Izaurralde E. mRNA quality control: an ancient machinery recognizes and degrades mRNAs with nonsense codons. FEBS Lett. 2007 Jun 19;581(15):2845–53.

34. Chess A. Mechanisms and consequences of widespread random monoallelic expression. Nat Rev Genet. 2012 May 15;13(6):421–8.

35. Meier JC, Kankowski S, Krestel H, Hetsch F. RNA Editing—Systemic Relevance and Clue to Disease Mechanisms? Front Mol Neurosci [Internet]. 2016 Nov 23 [cited 2019 Apr 10];9. Available from: https://www.ncbi.nlm.nih.gov/pmc/articles/PMC5120146/

