## Supplemental Material for "SmartRNASeqCaller: improving germline variant calling from RNAseq"

**Suppl figure 1: Difference in Precision and Recall and the baseline GATKpipeline**

Plot showing the difference in both precision and recall of the workflows compared with GATK best practices (in red). We can see how SmartRNASeqCaller allow a remarkable improvement in precision (+9.3% on the validation set), at a modest loss in recall (less than 1% on the validation samples) when applied on GATK results. SNPiR trades an increase in precision with a 20% loss in recall. . As noted before, the naive filtering schema is the worst method as it sacrifices most of the recall power with a modest improvement of precision.

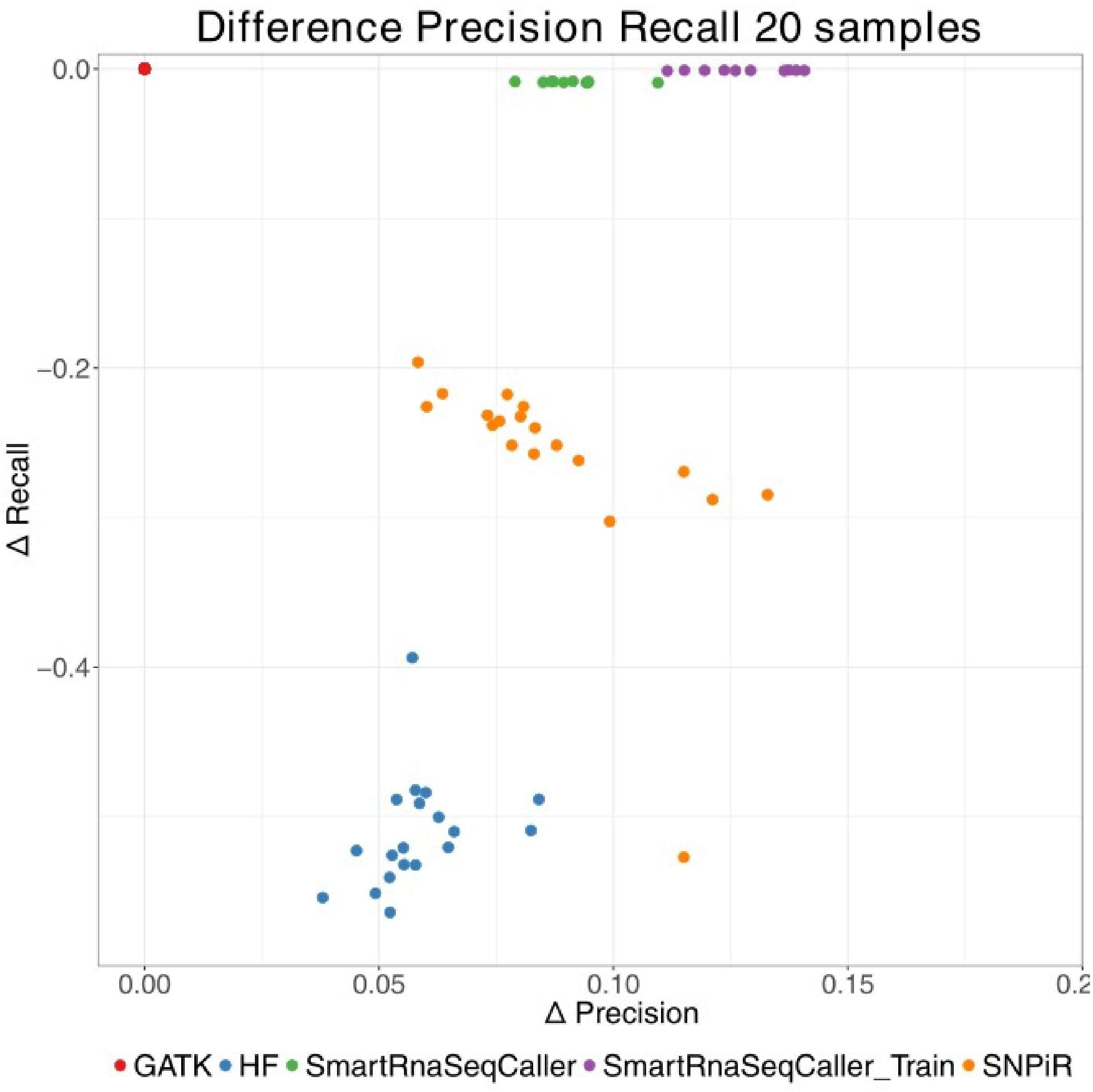
